# Aggregation of nontuberculous mycobacteria is regulated by carbon:nitrogen balance

**DOI:** 10.1101/631283

**Authors:** William H. DePas, Megan Bergkessel, Dianne K. Newman

## Abstract

Nontuberculous mycobacteria (NTM) are emerging opportunistic pathogens that form biofilms in environmental reservoirs such as household water systems and aggregate into phagocytosis-resistant clusters during infection. NTM constitutively aggregate *in vitro*, a phenotype typically considered to be a by-product of the mycolic-acid-rich cell wall. While culturing a model NTM, *Mycobacterium smegmatis*, in rich medium, we fortuitously discovered that planktonic cells accumulated in the culture after ∼3 days. By providing selective pressure for bacteria that disperse earlier, we isolated a strain with two mutations in the oligopeptide permease operon (*opp*). A mutant lacking the *opp* operon (Δ*opp*) dispersed earlier and more completely than wildtype (WT). We show that Δ*opp*’s aggregation defect was nutrient related; aggregation was restored by non-peptide carbon sources. Experiments with WT *M. smegmatis* revealed that growth as aggregates is favored when carbon is replete, while dispersal can be induced by carbon starvation. In addition, under conditions of low available carbon relative to available nitrogen, *M. smegmatis* grows as planktonic cells. By adjusting carbon and nitrogen sources in defined medium, we tuned the cellular C:N ratio such that *M. smegmatis* grows either as aggregates or planktonic cells. Lastly, we tested the effect of C:N balance on aggregation in clinically relevant NTM. Altogether, we show that NTM aggregation is a controlled process that is regulated by the relative availability of carbon and nitrogen for metabolism. Because NTM aggregation is correlated with increased virulence, these results may contribute to targeted anti-biofilm therapeutics.

**Importance:** Free-living bacteria can assemble into multicellular aggregates called biofilms. Biofilms help bacteria tolerate multiple stresses, including antibiotics and the host immune system. Differing environmental pressures have resulted in biofilm architecture and regulation varying among bacterial species and strains. Nontuberculous mycobacteria are a group of emerging opportunistic pathogens that utilize biofilms to adhere to household plumbing and showerheads and to avoid phagocytosis by host immune cells. Mycobacteria harbor a unique cell wall built chiefly of long chain mycolic acids that confers hydrophobicity and has been thought to cause constitutive aggregation in liquid media. Here we show that aggregation is instead a regulated process dictated by the balance of available carbon and nitrogen. Understanding that mycobacteria utilize metabolic cues to regulate the transition between planktonic and aggregated cells reveals an inroad to controlling aggregation through targeted therapeutics.

## Introduction

The adhesive biofilm matrix can serve as a physical barrier against external stresses such as desiccation and predation, can interact with and sequester antimicrobial agents, and can short-circuit phagocyte signaling (1–4). Additionally, the 3D structure of biofilms creates chemical gradients across a cellular population (5–9), resulting in a spectrum of physiologies and metabolisms which, along with genetic diversification and stochastic differences in gene expression, gives rise to substantial cell-to-cell heterogeneity (9–12). Heterogeneous bacterial communities demonstrate increased fitness compared to homogenous communities in a variety of models and experimental systems (11–13). Notably, most antibiotics target rapidly dividing bacteria, so slow-growing and dormant cells that develop in biofilms contribute to antibiotic tolerance (5, 10, 14–17).

Nontuberculous mycobacteria (NTM) are emerging pathogens that utilize biofilm formation for survival and persistence both in the host and in the non-host environment (18–22). NTM are adept at surviving standard water decontamination protocols and are commonly found in household water systems, often growing as biofilms (18, 23). NTM can infect healthy adults after repeated exposure and are especially dangerous to immunocompromised populations and patients with lung disorders such as Cystic Fibrosis (CF) and Chronic Obstructive Pulmonary Disease (COPD) (23–26). Infections with NTM can be very difficult to treat; *M. abscessus* lung infections, in particular, require long courses of antibiotic cocktails that have limited efficacy and extensive adverse side effects (24, 27, 28). The ability of *M. abscessus* to aggregate into cord-like aggregates correlates with increased pathogenicity in a zebrafish model and an enhanced ability to evade phagocytosis (19–21), indicating that the formation of multicellular structures by NTM is positively related to their sustained infection of hosts.

Bacteria have evolved to enter and exit from the biofilm state in response to species- and strain-specific environmental signals. Peculiarly, mycobacteria form *in vitro* biofilms in nearly all laboratory culture conditions; aggregating into hydrophobic clumps in shaking cultures and forming pellicle biofilms at the air/liquid interface of static cultures (29–32). While environmental parameters such as iron and CO_2_ affect mycobacterial pellicle maturation, cues driving the transition between planktonic cells and biofilms have not been identified due to the apparent absence of a true planktonic state (33, 34). Constitutive aggregation suggests either that mycobacteria express adhesive structures in response to signals that are very common in laboratory cultures, or that they have adapted to always grow as aggregates in aqueous environments. The latter possibility has become the dominant paradigm, exemplified by the common addition of detergents such as Tween 80 to mycobacterial cultures to prevent clumping (31, 32).

In this study, we set out to understand whether and how aggregation is regulated in NTM. Towards this end, we developed an assay to quantify mycobacterial aggregation in liquid media under varying nutritional environments. Contrary to the conventional wisdom, we found that aggregation and dispersal are regulated processes in a variety of NTM, both pathogenic and non-pathogenic, dictated in large part by the relative availability of carbon and nitrogen.

## Results

### Mycobacterial aggregates disperse as cultures age

During routine culture in a rich medium with no detergent, the model NTM *Mycobacterium smegmatis* MC^2^155 grows as aggregated clumps. However, we noted that non-aggregated (planktonic) cells accumulated after ∼40 hours of growth (Fig. 1A). We developed an assay to distinguish and quantify aggregated cells and planktonic cells over time. Briefly, culture replicates were harvested over time by passing an entire culture through a 10 μm cell strainer. The OD_600_ of cells that passed through the strainer (planktonic fraction) was immediately recorded. Aggregates that collected on the strainer were water bath sonicated in PBS + 24.8% Tween 20, and the OD_600_ of the resultant suspension was recorded (Fig. 1B). Phase contrast microscopy revealed that the planktonic fraction was composed mostly of single cells and small clusters (Fig. 1C). SEM of a representative aggregate revealed a densely packed structure (Fig. 1D). Performing this assay on *M. smegmatis* grown in rich medium + 0.2% glucose revealed a decrease in the aggregate fraction concurrent with planktonic cell accumulation after ∼40 hours of growth, suggesting a mechanism of controlled dispersal (Fig. 1E).

**Figure 1.**
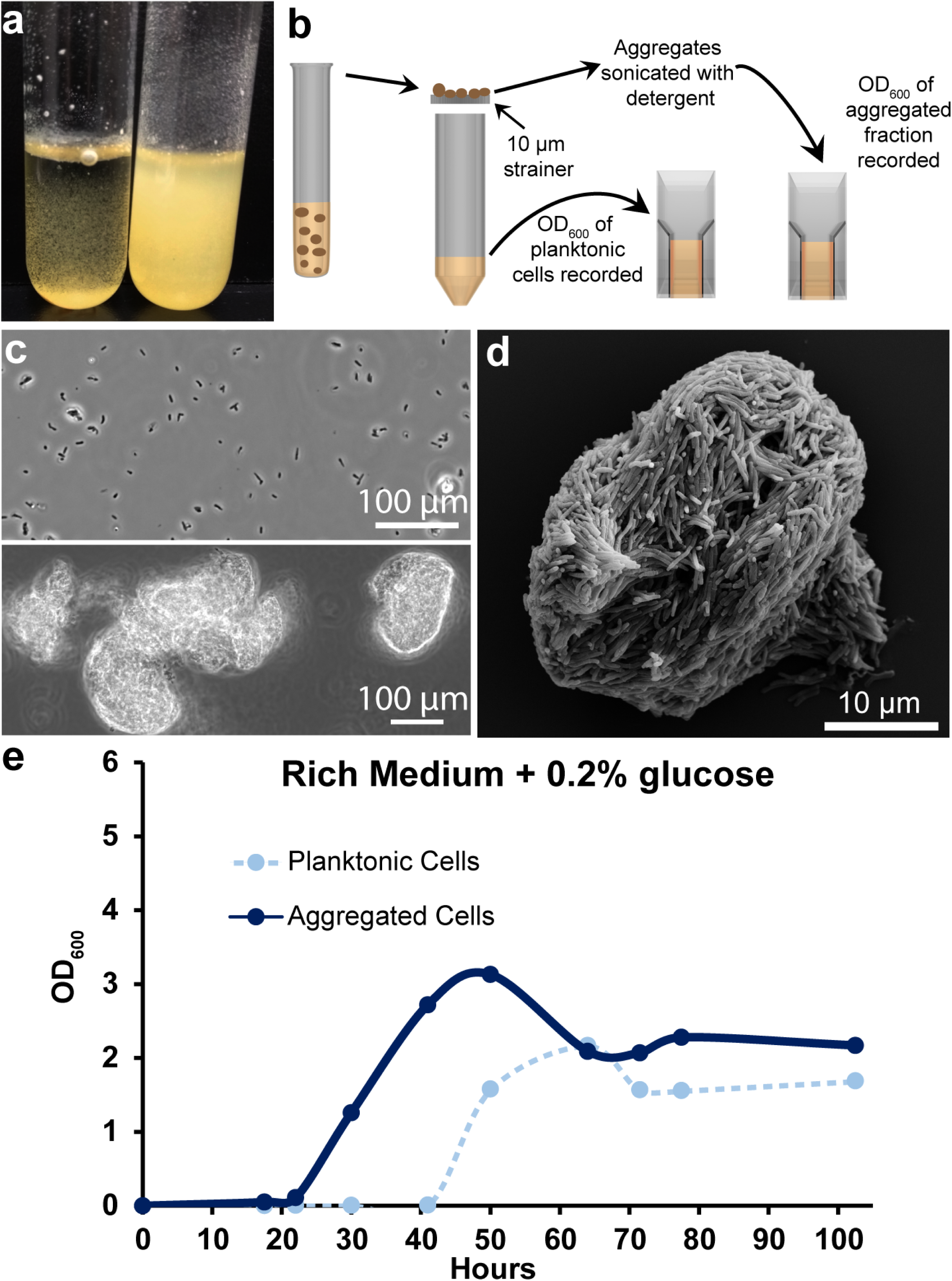
Quantification of mycobacterial aggregation/dispersal over time. (A) In rich medium + 0.2% glucose, *M. smegmatis* grows as clumps at early time points (left tube, 30 hours of growth). In older cultures, planktonic cells accumulate (right tube, 72 hours of growth). (B) Cartoon depicting a method to separate and quantify aggregated and planktonic mycobacterial cells. (C) Phase-contrast micrograph showing the planktonic (top panel) and aggregated (bottom panel) fraction of a 72-hour-old culture. The planktonic fraction is largely single cells and small clumps. Cells that are retained on the strainer (aggregated fraction) mostly exist as large clumps. (D) SEM of a representative *M. smegmatis* aggregate that was retained on the strainer after ∼30 hours of growth in rich medium. (E) Aggregation curve of WT *M. smegmatis* grown in rich medium + 0.2% glucose. Cells were harvested at each indicated timepoint and processed with the method outlined in Fig. 1B. Data are representative of n=4 trials.

### Mutations in oligopeptide permease genes cause early dispersal

To gain insight into the genetic regulation of *M. smegmatis* aggregation and dispersal, we designed an evolution experiment to select for mutants that disperse earlier than WT in rich medium + 0.2% glucose. Briefly, every 24 hours 1 mL of a 5 mL culture was centrifuged at low speed to pellet aggregates. A new 5 mL culture was inoculated with 100 μL of the supernatant and grown for another 24 hours (Fig. 2A). After 60 passages (roughly 575 doublings), planktonic cells visibly accumulated after 24 hours of growth. Passage 60 was plated and a single colony was selected and cultured. The passage 60 isolate displayed an early dispersal phenotype compared to WT in rich medium + 0.2% glucose (Fig. 2B). We sequenced the genomes of the passage 60 isolate, our WT strain (passage 0), and a passage 40 isolate that showed no early dispersal phenotype (Fig. S1). In total, the passage 40 isolate had 13 mutations compared to our passage 0 isolate, seven of which were in non-transposon open reading frames (ORFs). The passage 60 isolate had 11 mutations compared to our passage 0 isolate, nine of which were in non-transposon ORFs (Table 1).

**Figure 2.**
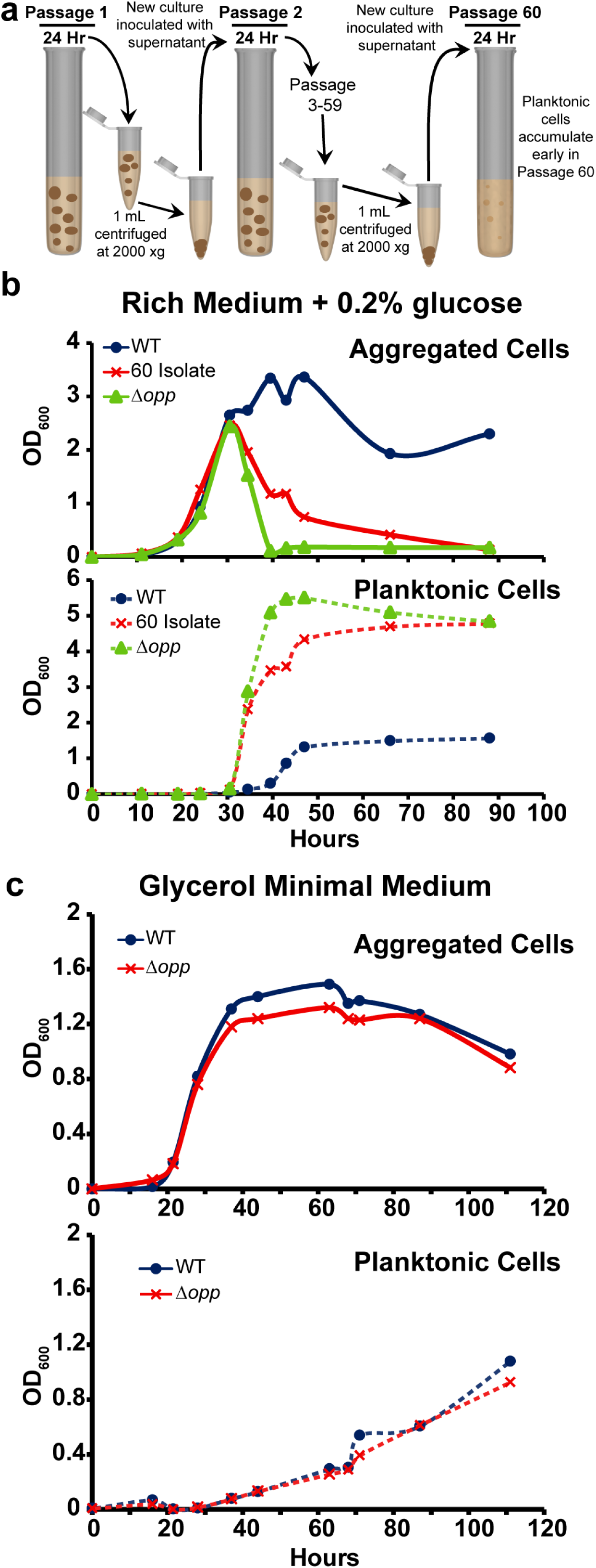
Mutations in an oligopeptide permease operon lead to early dispersal. (A) Cartoon depicting an evolution experiment to select for an *M. smegmatis* strain that disperses earlier than WT. (B) Aggregation curve of WT *M. smegmatis*, the passage 60 isolate, and Δ*opp* grown in rich medium + 0.2% glucose. The top panel shows the aggregated fraction and the bottom panel shows the planktonic fraction. Data are representative of n=3 trials. (C) Aggregation curve of WT *M. smegmatis* and Δ*opp* grown in glycerol defined medium. The top panel shows the aggregated fraction and the bottom panel shows the planktonic fraction. Data are representative of n=2 trials.

**Table 1.**

|  | Gene | Function | Mutation Type | Mutation |
| --- | --- | --- | --- | --- |
| Pass 40 | **MSMEG_1808** | SufE | **Missense** | **Glu39Val** |
|  | **MSMEG_2148** | HNH endonuclease domain-containing protein | **Frameshift** | **Ser534fs \*** |
|  | **MSMEG_5061** | Extracellular solute binding protein | **Missense** | **Ser249Pro** |
|  | **MSMEG_5808** | Binding protein dependent transporter | **Missense** | **Arg117Cys** |
|  | **MSMEG_6397** | Hypothetical protein | **Missense** | **Ser21Pro** |
|  | **MSMEG_6430** | Hypothetical protein | **Missense** | **Thr371Lys** |
|  | **MSMEG_6821** | NLP/P60 family protein | **Missense** | **Gln2017Arg** |
| Pass 60 | **MSMEG_0639** | Oligopeptide transport ATP-binding protein OppF | **Frameshift** | **Lys12fs #** |
|  | **MSMEG_0640** | Oligopeptide transport ATP-binding protein OppD | **Missense** | **Phe96Leu** |
|  | **MSMEG_2148** | HNH endonuclease domain-containing protein | **Missense** | **Pro380Arg** |
|  | **MSMEG_2148** | HNH endonuclease domain-containing protein | **Frameshift** | **Ser534fs \*** |
|  | **MSMEG_3677** | Serine/Threonine protein kinase | **Silent** | **Val320Val** |
|  | **MSMEG_4217** | DivIVA protein | **Missense** | **Glu107Gly** |
|  | **MSMEG_5061** | Extracellular solute binding protein | **Frameshift** | **Glu225fs &** |
|  | **MSMEG_5395** | Sensor Histidine Kinase KdpD | **Missense** | **Arg627Cys** |
|  | **MSMEG_6497** | Hypothetical protein | **Missense** | **His43Gln** |

Red text indicates the genes that were mutated in WT to test for aggregation defects
\* - MSMEG\_2148 is 544 amino acids. Ser534 frameshift hypothetically replaces the 8 C-terminal amino acids with a different 23 amino acid sequence.

To identify dispersal-related mutations, we narrowed our list of passage 60 candidate genes by discarding two genes that were also mutated in the passage 40 isolate (MSMEG_2148 and MSMEG_5061), one gene that acquired a silent mutation (MSMEG_3677), and *divIVA* (MSMEG_4217) because it is essential in *M. smegmatis* (35). We generated deletion mutants in a WT background of the four remaining candidates: *oppF*, *oppD*, *kdpD*, and the hypothetical gene MSEMG_6497. Because *oppF* and *oppD* code for two ATPase subunits associated with an oligopeptide permease (opp) complex, we deleted the entire 5-gene *opp* operon (MSMEG_0643-MSMEG_0639, termed Δ*opp*). While Δ*kdpD* and Δ*MSMEG_6497* showed no dispersal phenotype (Fig. S2), Δ*opp* phenocopied the passage 60 isolate by displaying early dispersal (Fig. 2B), indicating that a functional oligopeptide permease system helps maintain aggregation in rich medium + 0.2% glucose.

The Opp complex imports oligopeptides for signaling and/or catabolism in multiple bacterial species (36, 37). Our rich medium contains tryptone and yeast extract, both of which are composed largely of oligopeptides, so we reasoned that 1.) exogenous peptides themselves are a pro-aggregation signal, 2.) a self-produced peptide pheromone serves as a pro-aggregation signal, or 3.) metabolizing peptides as nutrients provides the cell with a pro-aggregation signal. To distinguish between these possibilities, we grew WT and Δ*opp* in a defined, peptide-free glycerol medium. If exogenous peptides are necessary for aggregation (1), neither WT nor Δ*opp* should aggregate in the peptide-free medium; if a self-produced pheromone is required for aggregation (2), WT should aggregate but Δ*opp* should be defective; if peptides are used as a nutrient that provides a pro-aggregation signal (3), providing the cells with alternative carbon and nitrogen sources should bypass the need for peptide import and both strains should aggregate. Both WT and Δ*opp* maintained aggregation to a similar degree in glycerol defined medium (Fig. 2C), suggesting that the Opp complex promotes aggregation in rich medium by increasing cells’ access to the peptide nutrient sources.

### Carbon availability dictates *M. smegmatis* aggregation and dispersal

Because Δ*opp*’s aggregation deficiency in rich medium + 0.2% glucose appeared to be due to a defect in nutrient uptake, we tested whether non-peptide carbon supplementation could complement this defect. Indeed, glucose addition prolonged aggregation in both WT and Δ*opp*, suggesting that carbon starvation is a signal for dispersal (Fig. 3A, Fig S3A). Because of the utility of being able to measure near-complete dispersal, rich medium experiments going forward contain no glucose unless otherwise noted. If carbon starvation leads to aggregate dispersal, we would predict that either carbon-free buffer or carbon-depleted medium should be sufficient to induce dispersal. We therefore resuspended WT aggregates (grown in rich medium for 48 hours) in either PBS or conditioned medium from 52-hour-old cultures. After 12 hours, we harvested and quantified aggregated and planktonic populations (Fig. 3B). Aggregates decreased to a similar degree in both conditioned medium and PBS (Fig. 3B). Furthermore, when 0.6% glucose was added to conditioned medium, dispersal was largely prevented (Fig. 3B). Unexpectedly, when aggregates were resuspended in conditioned medium, planktonic cells accumulated to a significantly higher extent compared to PBS (Fig. 3B). This result indicated that, instead of growth as aggregates and subsequent dispersal, there may be a window of time in a rich medium culture wherein nutrient conditions favor growth as planktonic cells.

**Figure 3.**
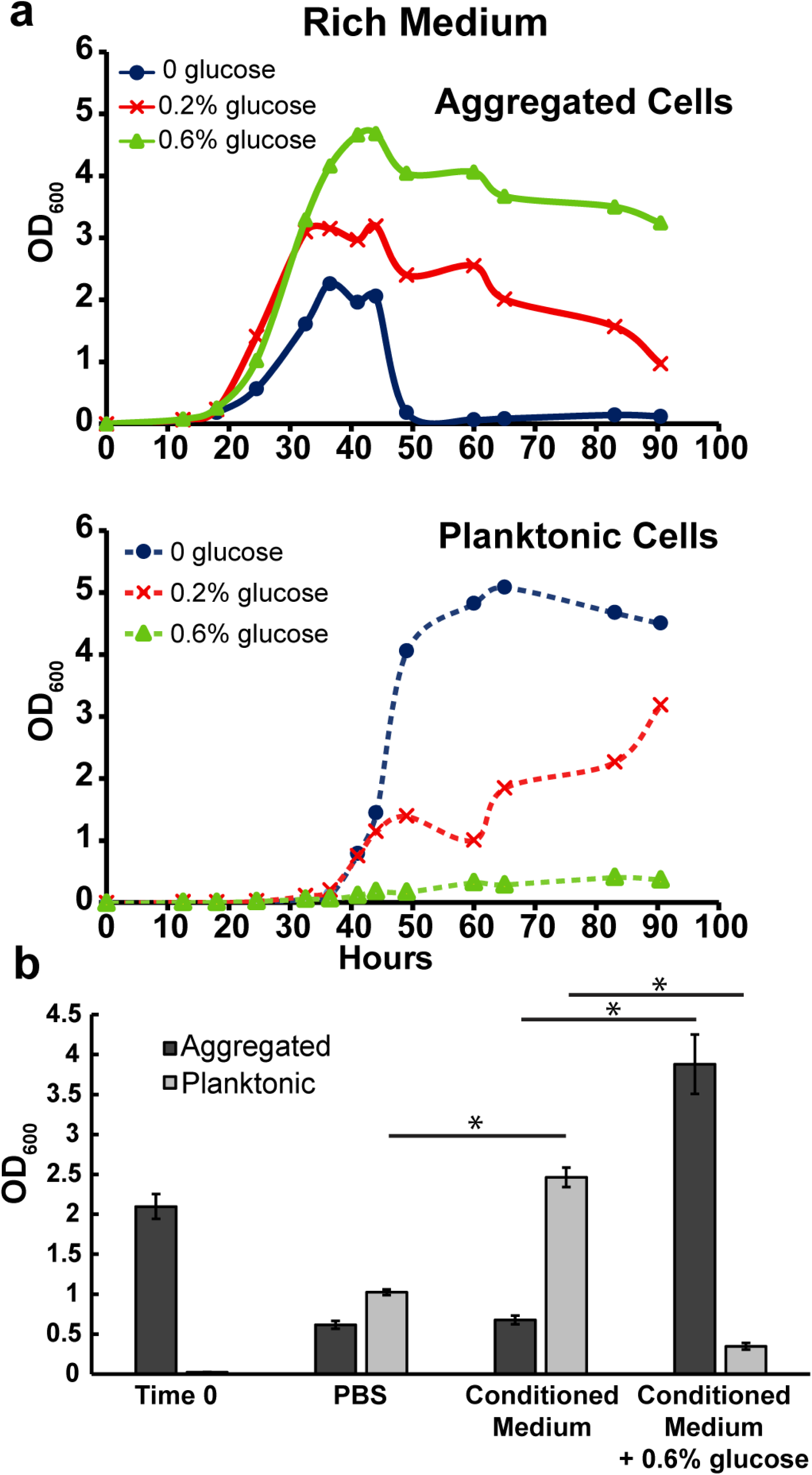
Carbon depletion leads to dispersal. (A) Aggregation curve of WT *M. smegmatis* in rich medium + no glucose, 0.2% glucose, or 0.6% glucose. The top panel shows the aggregated fraction and the bottom panel shows the planktonic fraction. Data are representative of n=3 trials. (B) Aggregates harvested from 48-hour-old rich medium cultures (Time 0) were resuspended in conditioned medium (filter-sterilized from 52-hour-old-rich medium cultures), PBS, or conditioned medium + 0.6% glucose and grown for 12 hours. Each bar is an average of biological triplicates and error bars represent standard deviation. Asterisks represents p <0.05 by the Student’s T-test.

### Low C:N ratio drives growth as planktonic cells

Because the OD_600_ has a limited range in which it can accurately measure cell density, we measured CFUs/mL of both aggregated and planktonic fractions over time in rich medium (Fig. 4A). This experiment revealed three distinct phases of growth. In phase I (∼0-40 hours), both fractions grow at similar rates with the aggregated fraction outnumbering the planktonic fraction by roughly 10 fold. In phase II (∼40-53 hours), planktonic cells continue growing while aggregated fraction growth ceases. Then, in phase III (at ∼53 hours onward), aggregates disperse and the planktonic fraction enters stationary phase (Fig. 4A). Our results from Fig. 3 suggest that carbon excess and depletion drive growth as aggregates and aggregate dispersal, respectively. Therefore, we sought to characterize the phase II culture conditions that favored planktonic cell growth. One well-characterized side effect of bacterial growth on peptides is the release of excess ammonium into the medium (38). Indeed, ammonium levels increased as our cultures aged, reaching ∼33 mM at 48 hours (Fig. 4B). To test whether ammonium facilitated growth as planktonic cells, we added excess NH_4_Cl to starting cultures and tracked aggregation. Ammonium addition led to earlier accumulation of planktonic cells and reduced aggregation (Fig. 4C, S3). To test whether salts have a general effect on aggregation, we added 75 mM NaCl to WT cultures. NaCl did not affect aggregation kinetics, indicating that ammonium specifically favors planktonic growth (Fig. S4). If the high ammonium concentration in conditioned medium favors growth as planktonic cells, it is notable that adding excess carbon to conditioned medium shifts the population back towards growth as aggregates (Fig. 3B). Altogether, these results are consistent with a model wherein carbon replete conditions favor growth as aggregates, high nitrogen (relative to carbon) conditions favor growth as planktonic cells, and carbon depletion leads to aggregate dispersal.

**Figure 4.**
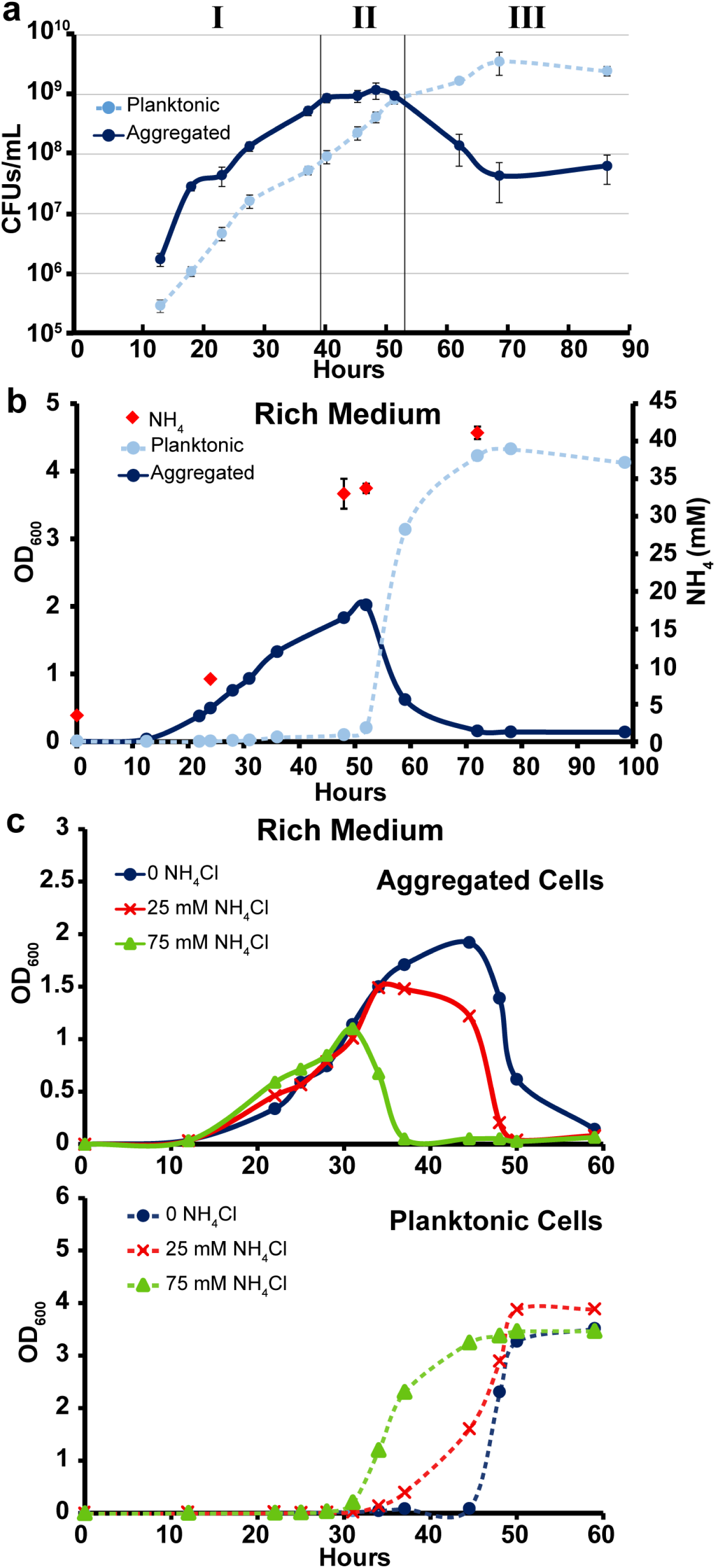
Low C:N availability favors growth as planktonic cells. A.) CFUs/mL for WT *M. smegmatis* grown in rich medium (no glucose). Each data point is the average of biological triplicates and error bars represent standard deviation. Roman numerals denote three phases of growth as described in text. B.) Aggregation curve of WT *M. smegmatis* in rich medium (no glucose). At indicated time points, three additional cultures were harvested for NH_4_ IC measurements. Each NH_4_ data point is an average of biological triplicates and error bars represent standard deviation. Aggregation curve data are representative of n=5 trials. C.) Aggregation curve of WT *M. smegmatis* in rich medium (no glucose) with no NH_4_Cl, 25 mM NH_4_Cl, or 75 mM NH_4_Cl. The top panel shows the aggregated fraction and the bottom panel shows the planktonic fraction. Data are representative of n=3 trials.

### Defined medium designed for growth as aggregated or planktonic cells

To test whether *M. smegmatis* is able to grow as planktonic cells at low C:N ratios, we designed defined medium to supply the bacteria with either replete carbon and low nitrogen (high C:N) or replete nitrogen and low carbon (low C:N). To grow *M. smegmatis* with high C:N availability, we used glycerol as the main carbon source, glutamate as the main nitrogen source, and no ammonium (117 mM carbon, 5.5 mM nitrogen, C:N of the medium = 21.4). Glycerol is commonly supplied to mycobacteria because it supports fast growth and, as a small (three-carbon) uncharged molecule, can presumably passively diffuse across the mycolic acid barrier (39). Indeed, growth on glycerol floods most central metabolite pools compared to growth on other carbon sources in *Mycobacterium tuberculosis* (40). To generate low C:N availability, we used a charged three-carbon compound, pyruvate, as the main carbon source, and added 20 mM NH_4_Cl in addition to glutamate as the nitrogen source (117 mM carbon, 25.5 mM nitrogen, C:N of the medium = 4.58). In some bacteria, the relative availability of carbon and nitrogen sources can be reflected in total C:N content of the cell (41). Therefore, to assess whether our medium was effectively providing high or low C:N availability, we directly measured the ratio of cellular carbon to cellular nitrogen (by mass) of *M. smegmatis* grown in either pyruvate or glycerol medium when the total OD_600_ was between 0.5 and 0.7. As predicted, *M. smegmatis* grown on glycerol had a C:N ratio of 6.95, (stdev 0.85), and on pyruvate + NH_4_Cl had a C:N ratio of 5.02 (stdev 0.31, p = 0.005). Consistent with our hypothesis, *M. smegmatis* grew mostly as aggregates on glycerol and grew mostly as planktonic cells on pyruvate (Fig. 5A,B).

**Figure 5.**
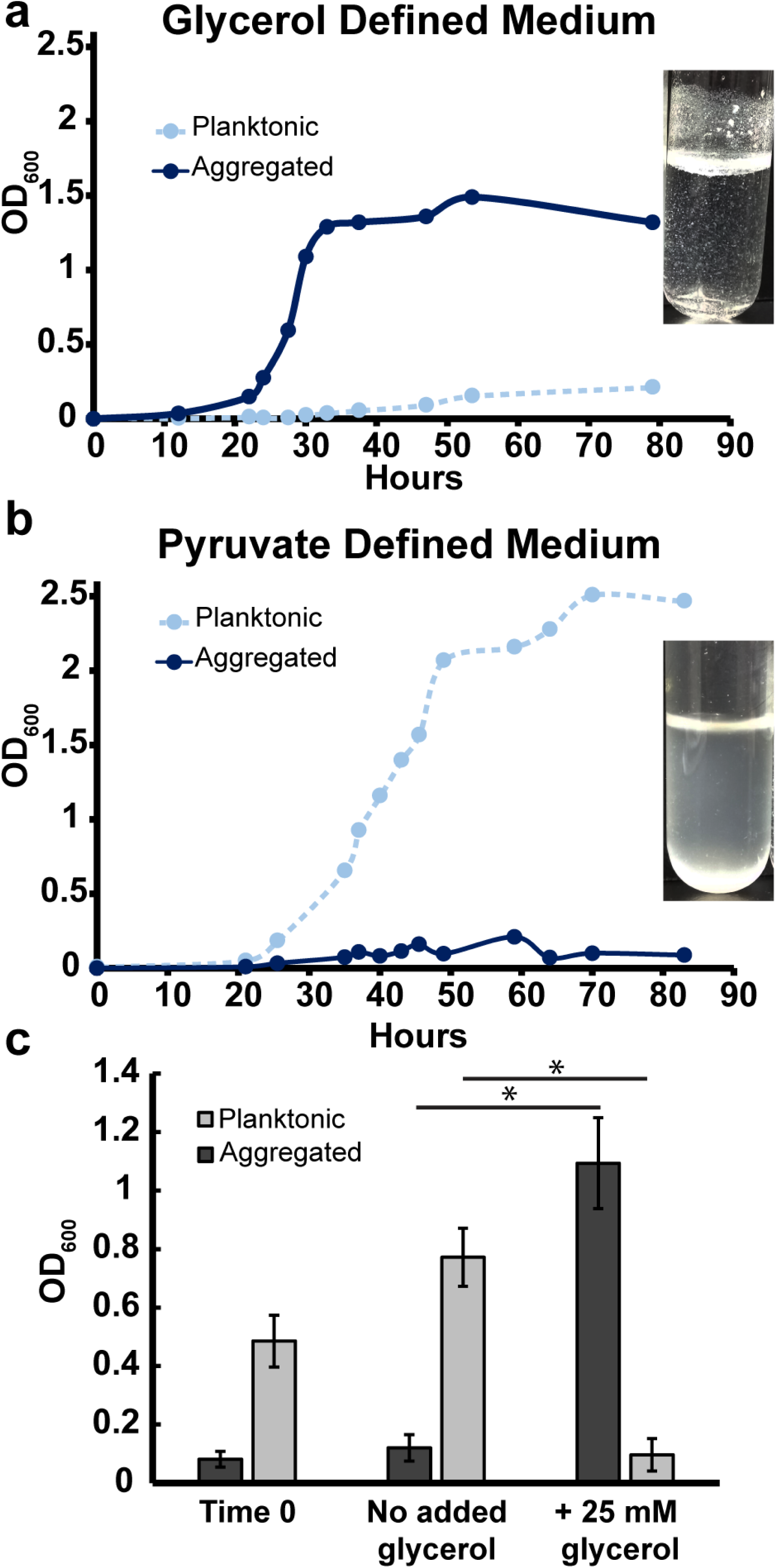
Defined medium designed to favor growth as aggregates or planktonic cells. (A) Aggregation curve of WT *M. smegmatis* in glycerol defined medium. Culture image was taken after 27 hours of growth. Data are representative of n=4 trials. (B) Aggregation curve of WT *M. smegmatis* in pyruvate defined medium. Culture image was taken after 34 hours of growth. Data are representative of n=4 trials. (C) WT *M. smegmatis* was grown in pyruvate + NH_4_Cl minimal medium for 34 hours (Time 0). Glycerol was then added to 25 mM, and cultures were incubated for six more hours before harvesting. Bars represent biological triplicates and error bars represent standard deviation. Asterisks represents p <0.05 by the Student’s T-test.

The ratio of C:N in natural environments such as soil affects bacterial diversity and growth and is often tuned in order to favor desired bacterial metabolisms in industrial settings (42, 43). It is therefore notable that even when grown in pyruvate defined medium with no ammonium (117 mM carbon, 5.5 mM nitrogen, C:N of the medium = 21.4, equal to glycerol defined medium), *M. smegmatis* had a relatively low cellular C:N ratio of 5.23 (stdev 0.38, p = 0.01 compared to glycerol grown cells) and grew as mostly planktonic cells (Fig. S5). These results reinforce that the form of available nutrients, and not just total carbon and nitrogen in an environment, can impact a cell’s C:N status and dependent phenotypes.

Lastly, we leveraged our pyruvate defined medium to test whether planktonic cells can transition to aggregates. Planktonic *M. smegmatis* was grown for 36 hours in pyruvate + NH_4_Cl defined medium before addition of 0 or 25 mM glycerol. By six hours post glycerol addition, the majority of the planktonic population had aggregated (Fig. 5C), further demonstrating that aggregation state is dynamic and dependent on the ratio of available C:N.

### C:N-dependent aggregation regulation is common among NTM

To determine whether C:N regulation of aggregation is conserved among clinically relevant NTM, we grew type strains of *M. abscessus* and *M. fortuitum* along with four *M. abscessus* subsp. *abscessus* clinical isolates (two rough colony isolates and two smooth colony isolates) in rich medium and tracked aggregation kinetics (Fig. 6, S6). Both type strains and one smooth colony clinical isolate accumulated planktonic cells at later culture timepoints, with glucose addition increasing total aggregation and ammonium addition favoring growth as planktonic cells (Fig. 6, S6). Neither rough colony *M. abscessus* isolate accumulated planktonic cells, even after addition of ammonium. In contrast, the smooth colony isolate that did not disperse in rich medium grew solely as planktonic cells when provided with supplemental ammonium. Altogether, our results indicate that C:N balance is a common regulator of NTM aggregation, with rough colony clinical isolates being a possible exception.

**Figure 6.**
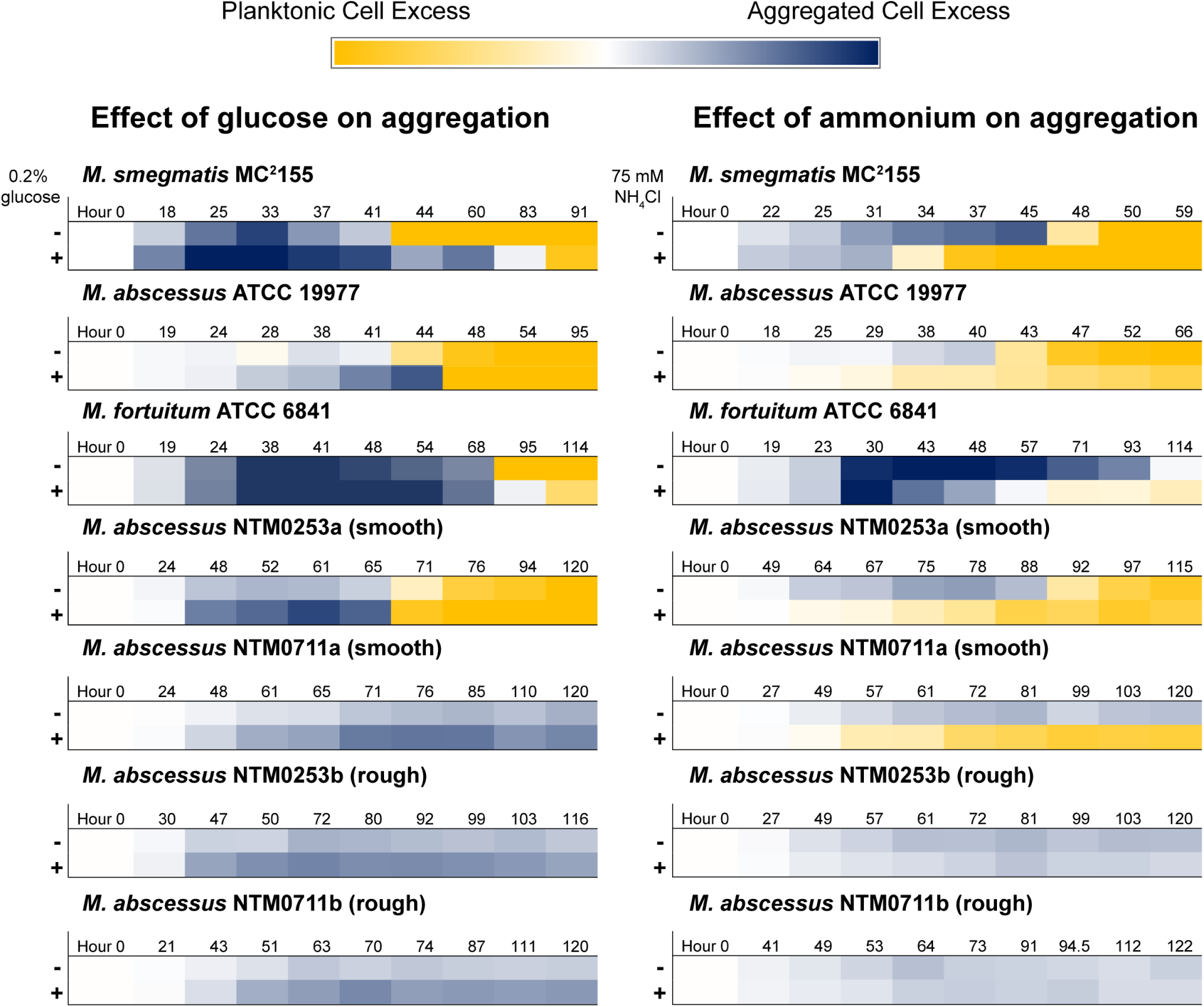
C:N regulation of aggregation/dispersal is common among NTM. Aggregation curves in rich medium +/- 0.2% glucose (left column) or rich medium +/- 75 mM NH_4_Cl (right column) were recorded for indicated strains. Ten timepoints were selected from each curve to span the entire timecourse. The OD_600_ value of the planktonic fraction was multiplied by −1, and then the OD_600_ values of both fractions were added together. The darkest blue color corresponds to sums of 2.5 or greater and the darkest yellow color corresponds to sums of −2.5 or less. Times are rounded up to the nearest hour. Data are representative of at least n=2 trials. The *M. smegmatis* heatmaps represent aggregation curves shown in Fig. 3A and Fig. 4C. The aggregation curves from which the other heatmaps were derived are included in Fig. S6.

## Discussion

The role of biofilm formation in rendering bacteria recalcitrant to antibiotics and immune killing provides motivation to develop novel anti-biofilm strategies. However, because bacteria have evolved to occupy and form biofilms in diverse ecological niches, the regulatory pathways and physical components that govern biofilm formation differ significantly between species. As such, a species-specific, in-depth understanding of how cells sense and respond to their environment by aggregating under certain conditions, and growing as planktonic cells under others, is essential in order to control bacterial biofilm formation for any specific pathogen. In this work, we have found a role for C:N balance in dictating the transition between planktonic and aggregated states in NTM.

Understanding the environmental niches in which NTM have evolved can lend context to our finding that C:N balance controls aggregation state. NTM are non-motile saprophytes that are common residents of soil and waterways (22, 28, 44). In soil, carbon is most often the limiting nutrient for bacterial growth (45, 46). At a low C:N ratio, our data suggest that NTM could exist at least partly as planktonic cells. As water flow is a major factor in determining movement of bacteria through soil (47, 48), NTM in this state might be susceptible to water-mediated transport to another region of the rhizosphere (potentially containing more carbon). Larger bacterial cell sizes correlate with decreased movement through soil (49). Therefore, if NTM were growing as aggregates in carbon-rich conditions, we would expect them to be less likely to be washed away into potentially more carbon-depleted regions. While speculative, this natural ecological context motivates us to consider how mycobacteria might sense the C:N balance in their environment and control their aggregation state accordingly.

It is well appreciated that carbon and nitrogen availability dictate the metabolic and growth capacity of a cell (50, 51), and bacteria are able to coordinate carbon and nitrogen metabolism through a variety of means (52). The cellular C:N ratio provides a rough estimate of the cell’s C:N status, but it is not a parameter that a cell can directly sense. How then do mycobacteria translate C:N availability to aggregation? Our data show that no one carbon source is necessary to drive aggregation. Interestingly, by responding to flux through a metabolic pathway, a cell can integrate the signal from multiple inputs without needing to measure each one specifically (53). It thus seems possible that mycobacteria sense and respond to flux-dependent metabolites – molecules whose intracellular pools correlate with flux through specific metabolic pathways, such as fructose-1,6-bisphosphate (FBP), the levels of which correlate with glycolytic flux (53, 54), or 2-oxoglutarate (2OG), the levels of which correlate to flux through the TCA cycle (53, 55). Alternatively, or in addition, two-component systems might mediate the translation between metabolite availability and aggregation. Uncovering the pathways through which NTM achieve aggregation control is a priority for future work.

Regardless of the signal transduction mechanism, a surface adhesin must mediate the aggregation phenotype. Like many members of the *Corynebacteriales* order, mycobacteria produce a mycomembrane: a cell wall composed of peptidoglycan, arabinogalactan covalently linked to an inner leaflet of long-chain mycolic acids, and an outer layer of extractable lipids, lipoglycans, and proteins (56, 57). As such, the mycobacterial cell wall is unusually lipid rich (58, 59). A lipid-rich cell wall fits the long-standing observation that mycobacteria clump together into hydrophobic aggregates; in his original description of *M. tuberculosis* in 1882, Robert Koch noted that the bacteria “…ordinarily form small groups of cells which are pressed together and arranged in bundles” (60). Clumping (or cording, depending on aggregate morphology) is now recognized as a ubiquitous feature of pathogenic and nonpathogenic mycobacteria (21, 32, 61). As clumps are hydrophobic, detergents such as Tween 80 are almost universally added to mycobacterial cultures to favor growth as dispersed cells (32, 61, 62).

Inherent to the chemical intuition linking a lipid-rich cell wall and spontaneous clumping is the assumption that mycobacteria display a *constitutively* hydrophobic cell surface. Several studies of the mycomembrane composition challenge this dogma. Trehalose dimycolate (TDM) was originally called ‘cord factor’ because cording is reduced when TDM is removed from the cell envelope via petroleum ether extraction (63–65). TDM expression is regulated by sugar availability in *M. avium*, implying that TDM-mediated aggregation can be controlled by the cell in response to the environment (66). Likewise, mycolic acid chain length affects aggregation (67), and *M. smegmatis* can regulate mycolic acid chain length in response to environmental factors (29, 68). Finally, genes involved in the biosynthesis and glycosylation of cell-surface glycopeptidolipids (GPLs) in *M. smegmatis*, *M. avium*, and *M. abscessus* affect aggregation and cell surface hydrophobicity (69–72). GPL production and glycosylation are also regulated by chemical signals (70, 73). In addition to providing evidence that mycobacteria can dynamically regulate cell envelope composition and surface hydrophobicity, these studies provide candidate adhesins that could be effectors of C:N-driven aggregation regulation.

Finally, the fact that NTM regulate aggregation has potentially important biomedical relevance. New treatments are needed to combat NTM infections, such as those caused by *M. abscessus,* which is notoriously difficult to eradicate. It is noteworthy that that rough colony isolates of *M. abscessus* subsp. *abscessus* do not disperse in rich medium. Rough *M. abscessus* isolates are typically the result of mutations that reduce GPL production (72, 74, 75). Accordingly, we might hypothesize that C:N regulation is linked to GPL production or modification, which directly impacts the aggregation state. It is worth exploring whether nodes along such a pathway could be identified and exploited as new targets for biofilm control. The rising threat of NTM infections, particularly to susceptible communities such as CF patients, as well as the correlation between increased aggregation and virulence, lends motivation to further probe the mechanisms of aggregation and dispersal in these pathogens (19, 21, 27, 28).

## Materials and Methods

### Strains and growth conditions

Strains, plasmids, and primers used in this study are listed in table S1. The rich medium used in this study was TYEM (10 g tryptone, 5 g yeast extract/L + 2 mM MgSO_4_). Where noted, filter sterilized glucose or NH_4_Cl were added as supplements to autoclaved TYEM. For routine culturing of mycobacteria, bacteria were grown in TYEM for ∼50-70 hours, at which time cultures were passed through 10 µm strainers (from pluriSelect, 43-50010-03) and planktonic cells were collected and processed. The exception was rough *M. abscessus* isolates NTM0253b and NTM0711b, which were cultured in TYEM + 0.05% Tween 80. Our defined medium was modified M63 -- 13.6 g KH_2_PO_4_ was dissolved in 500 mL Nanopure H2O and the pH was adjusted to 7.0 via addition of KOH. This 2X stock was filter sterilized and diluted to 1X with Nanopure H_2_O while adding filter sterilized supplements: MgSO_4_ to 1 mM, FeSO_4_ to 10 uM, SL-10 trace metal solution to 1x, proline to 0.5 mM, sodium glutamate to 5 mM, NH_4_Cl to 20 mM (when noted), and either glycerol to 30 mM or sodium pyruvate to 30 mM. Mutants in *M. smegmatis* MC^2^155 were made via recombineering as described with minor alterations (76). Briefly, *M. smegmatis* transformed with pJV53 was grown in TYEM + 0.05% Tween 80 + 25 ug/mL kanamycin until it reached an OD_600_ of 0.4-0.5. Acetamide was added to 0.2% and cells were incubated for 3 hours shaking at 250 rpm at 37°C. Cells were then made electrocompetent by serial washes with chilled 10% glycerol (1/2, 1/10th, 1/20th, 1/100^th^ original volume) with centrifugation at 4000 xg for 10 minutes at 4°C between washes. 100 uL of the cell mixture was then electroporated with 200 ng of linear DNA encoding a gentamicin resistance cassette (PCR amplified from plasmid pMQ30) flanked by 400-500 bps of sequence upstream and downstream of the target genes. Flanking regions were PCR-amplified from WT *M. smegmatis* colonies and Gibson assembly was utilized to combine flanking regions with the gentamicin resistance cassette. After mutagenesis, mutant strains were cured of pJV53 by passaging on TYEM with no antibiotics 3-7 times until they were verified as kanamycin-sensitive.

### Light microscopy and SEM

For light microscopy, samples were loaded onto Tekdon poly-L-lysine coated slides and phase contrast images were acquired on a Zeiss AxioObserver.A1 using a 40x 1.3 NA oil immersion objective. For SEM, WT *M. smegmatis* was grown in rich medium for 24 hours, at which point the culture was passed through a 10 µm strainer and washed with PBS. Aggregates that collected on the strainer were fixed in 4% PFA for 2 hours at room temperature, washed 2x with PBS, and fixed in 1% OsO_4_ for 1 hour at room temperature. After two more rinses with PBS, aggregates were dehydrated in an ethanol series, with 10 minute incubations in 50%, 70%, 90%, 95%, 100% ethanol, and a final incubation in 100% ethanol for 1 hour. Samples were then incubated in a 1:2 solution of hexamethyldisilazane (HMDS):ethanol for 20 minutes, a 2:1 solution of HMDS:ethanol for 20 minutes, followed by two incubations in 100% HMDS for 20 minutes each. Samples were then loaded onto silicon wafers, air dried, and attached to imaging stubs with conductive tape. Samples were sputter coated with 10 nm of palladium and imaged on a Zeiss 1550VP field emission SEM using an SE2 detector.

### Aggregation assays

Medium for aggregation assays was prepared in flasks and inoculated with the indicated strain of bacteria to an OD_600_ of 0.01. After mixing, 5 mL aliquots were pipetted into brand-new borosilicate disposable culture tubes. These culture replicates were incubated at 37°C while shaking at 250 rpm. At indicated timepoints, a single culture replicate was harvested by pouring the entire culture through a 10 µm strainer. Culture that passed through the strainer was designated as the planktonic cell fraction and the OD_600_ was immediately recorded. The original culture tube was washed with 5 mL of PBS, which was then poured over the aggregate fraction to remove residual planktonic cells. Aggregates that remained on the strainer were resuspended in 4.5 mL PBS + 6% Tween 20 and poured back in the original culture tube. 500 µL of Tween 20 was added for a final volume of 5 mLs and a final Tween 20 concentration of 28.5%. Aggregate fractions were then water bath sonicated until no visible clumps remained, and the OD_600_ of the aggregate fraction was recorded. For CFU counts, a slightly modified protocol was employed for the aggregate fraction. Instead of PBS, aggregates were resuspended in TYEM + 0.05% Tween 80, to which 100 µL of autoclave-sterilized Tween 20 was added. Aggregates were then water bath sonicated until no clumps were visible. Both planktonic and aggregate fractions were then serially diluted in TYEM + 0.05% Tween 80 and serial dilutions spanning seven orders of magnitude were plated on TYEM agar plates as 10 µL drips. Plates were incubated at 37°C for ∼2 days and colonies were counted at the appropriate dilution. Conditioned medium was prepared by centrifuging 52-hour-old cultures and filtering the supernatant through a 0.2 µm filter. For conditioned medium experiments, three 48-hour-old cultures were pooled by passing them through a single 10 µm strainer. Aggregates were washed with 5 mL of PBS and then resuspended in 15 mL of conditioned medium (or PBS as indicated). 5 mL aliquots were partitioned into three culture tubes, and after 12 hours of shaking at 37°C, aggregates and planktonic cells were separated and quantified.

### Evolution Experiment/Sequencing

WT *M. smegmatis* was inoculated into TYEM + 0.2% glucose. After 24 hours, 1 mL of culture was centrifuged for 1 minute at 2000 x g. 100 µl of supernatant was inoculated into a new TYEM + 0.2% glucose culture. The process was repeated every 24 hours. After 60 passages, planktonic cells were visibly accumulating at 24 hours. This culture was plated on TYEM agar plates and a single colony was selected as the passage 60 isolate. Along with an isolate from passage 0 and passage 40, this strain was grown to mid-exponential phase and DNA was extracted as described (77). DNA was fragmented using the NEBNext dsDNA Fragmentase (New England Biolabs, Ipswich MA) according to the manufacturer’s instructions. Briefly, 1 µg of passage 0 and passage 40 DNA and 725 ng of passage 60 DNA were treated with fragmentase for 15 minutes in order to achieve an acceptable size distribution, which was assessed using a High-Sensitivity DNA chip on a Bioanalyzer instrument (Agilent). Libraries for sequencing were prepared using the NEBNext DNA Library Prep kit according to instructions, which included end-repair of the fragments, dA-tailing, and ligation to adaptors. Each sample was PCR-amplified with a universal primer and a unique bar-coded primer, using 12 amplification cycles. Final libraries were verified using a Bioanalyzer High-Sensitivity DNA chip and quantified using the Qubit fluorimeter and dsDNA dye (Invitrogen). Sequencing was performed by the Millard and Muriel Jacobs Genetics and Genomics Laboratory at the California Institute of Technology using the Illumina HiSeq 2500 platform. Approximately 15 million single reads of 50 bp each were collected for each sample. Base-calling and de-multiplexing were performed by the Illumina HiSeq Control Software (HCS, version 2.0). The resulting FASTQ files were concatenated into one file per sample and filtered and trimmed by quality score per base using the Trimmomatic software package with the following parameters: LEADING:27 TRAILING:27 SLIDINGWINDOW:4:20 MINLEN:35 (78). Surviving reads were mapped to the *Mycobacterium smegmatis* str. MC2 155 genome (gi|118467340|ref|NC_008596.1) using bwa (version 0.7.12) (79), and sorted and converted to binary format using SAMtools (version) (80). Tools from the Genome Analysis Tool Kit (GATK, version 2.7-4-g6f46d11) (81) were used to call SNPs and small insertions and deletions relative to the reference genome as follows: first, duplicate reads were identified and marked using the MarkDuplicates tool. Next, putative insertions and deletions were identified using the RealignerTargetCreator tool, and reads surrounding them were re-aligned using the IndelRealigner tool. Finally, putative variants relative to the reference genome were called using the UnifiedGenotyper tool. 144 variant regions were confidently identified in the passage 0 sample, 153 variant regions were identified in the passage 40 sample, and 154 variant regions were identified in the passage 60 sample. Most of these variations were common to all three samples and were not considered further. For mutations of interest, the effects on protein coding sequence were predicted using the snpEff tool (version SnpEff 4.3t) (82). Genes affected by variations in non-transposon ORFs arising in the passage 40 and passage 60 sample relative to the passage 0 sample are listed in table 1.

### Ammonium measurements

At the time points indicated, 1 mL of culture was centrifuged at 16,000 × *g* at room temperature for 1 minute to pellet cells. Supernatants were filter sterilized through a 0.2 µm syringe filter and diluted 1:40 in nanopure H_2_O. Parallel ion chromatography systems operated simultaneously (Dionex ICS 200, Environmental Analysis Center, Caltech) were used to measure ammonium. A single autosampler (Dionex AS 40) loaded both systems’ sample loops serially. The 5 µL sample loop on the anion IC system was loaded first, followed by a 5 µL sample loop on the cation IC system. Both columns and both detectors were maintained at 30°C. Anionic components in the sample were resolved using a AS-19 separator (2×250mm) column protected by an AG-19 guard (2×50mm). A hydroxide gradient was produced using a potassium hydroxide eluent generator cartridge and pumped at 0.25 mL per minute. The gradient began with a 10 mM hold for 10 minutes, increased linearly to 58 mM at 25 minutes, remaining at 58 mM until the end of data acquisition at 32 minutes. Seven minutes were allowed between analyses to return the column to initial conditions. Anions were detected at neutral pH using an AERS - 500 2mm suppressor (Thermo) operated in eluent recycle mode with an applied current of 30 mA and conductivity detection cell maintained at 35°C. A carbonate removal device (CRD 200 2mm) was installed between the suppressor eluent out and the conductivity detector eluent in ports. Ammonium, calcium, magnesium, potassium and sodium were resolved using a CS-12A separator column (2×250mm) protected by a CG-12A guard column (2×50). Isocratic methylsulfonate at 20 mM was produced using a methylsulfonic acid based eluent generated cartridge and pumped at 0.25 mL per minute. Suppressed conductivity detection using a Dionex CERS-500 2 mm suppressor operated in eluent recycle mode with an applied current of 15 mA. Ammonium standards ranging from 1 µM to 1 mM (1 µM, 10 µM, 50 µM, 100 µM, 500 µM, and 1 mM) were run along with samples. A standard curve was generated by fitting a quadratic curve to standard measurements.

### C:N measurements

For defined medium conditions, 16 5 mL cultures (either in pyruvate defined medium +/- NH_4_Cl or glycerol defined medium) were grown to an OD_600_ between 0.5 and 0.7. The 16 cultures were divided into two sets of eight cultures. All eight cultures in a set were poured into a single 50 mL conical tube. Samples were then centrifuged at 6000 x g for 10 minutes at 4°C. Pellets were then washed 2x with 25 mL PBS, with centrifugation in between. After the second wash, each pellet was resuspended in 1.2 mL PBS, which was divided among two 1.5 mL centrifuge tubes in 600 µL aliquots (for a total of four samples/condition). After centrifugation at 16000 x g for 1 minute, supernatants were pipetted off and pellets were flash frozen in liquid nitrogen and stored at −80°C. Frozen samples were lyophilized, and ∼50 µg (for carbon measurement) and ∼700 µg (for nitrogen measurement) of each sample was weighed into an OEA lab tin capsule (pressed, ultra-clean, C61480.096P). Carbon and nitrogen were measured separately due to differing sensitivities of the instrument. Each sample was combusted in a Thermo Fisher EA IsoLink combustion system by oxidation at 1020°C over tungstic oxide, followed by reduction over elemental copper packed in the same furnace. The generated CO_2_ and N_2_ carried by a continuous helium flow (100ml/min) were subsequently passed through a water trap and then a 5Å molecular sieve GC at 50°C. The GC was used to separate N_2_ from CO_2_. Carbon and nitrogen were then diluted with helium in a Conflo IV interface/open split prior to entering the Thermo Fisher Delta V IRMS system for analysis. Depending on the configurations of the IRMS, either CO_2_ or N_2_ was measured for its total abundance. Integrated peak areas for both CO_2_ and N_2_ were calibrated by running urea standards, and empty tins were included as blanks. A Student’s T-test was used to generate p-values comparing conditions.

## Acknowledgements

We thank the Cystic Fibrosis Foundation (grants DEPAS17F0 to W.H.D. and BERGKE16F0 to M.B.) and the N.I.H (grant 1R01AI127850-01A1 to D.K.N.) for supporting our research. Lindsay Caverly (CS Mott Children’s Hospital, University of Michigan, Ann Arbor, MI, USA) graciously provided *M. abscessus* clinical isolates (NTM0253a, NTM0253b, NTM0711a, and NTM0711b), William Jacobs (Albert Einstein College of Medicine, Bronx, NY, USA) provided WT *M. smegmatis* MC^2^155, and William Bishai (Johns Hopkins University School of Medicine, Baltimore, MD, USA) provided plasmid pJV53. Igor Antoshechkin and the Millard and Muriel Jacobs Genetics and Genomics Laboratory at Caltech assisted with genome sequencing. Alex Sessions and Fenfang Wu (Caltech) helped with C:N measurements and analysis. Ion chromatography instrumentation used for this work is located in the Environmental Analysis Center at Caltech. We acknowledge Nathan F. Dalleska, Lev Tsypin, and Melanie Spero for Ion Chromatography method support. SEM was performed at the Caltech GPS Division Analytical Facility with the assistance of Chi Ma.

---

**Figure S1 – Passage 40 isolate displays no aggregation defect**

Aggregation curve of passage 0 (WT), 40, and 60 isolate in rich medium + 0.2% glucose. The top panel shows the aggregated fraction and the bottom panel shows the planktonic fraction. Data are representative of n=2 trials.

**Figure S2 – Neither Δ*kdpD* (MSMEG_5395) nor Δ*MSMEG_6497* have aggregation defects**

Aggregation curve of WT, Δ*kdpD* (MSMEG_5395), and Δ*MSMEG_6497* in rich medium + 0.2% glucose. The top panel shows the aggregated fraction and the bottom panel shows the planktonic fraction. Data are representative of n=2 trials.

**Figure S3 – Response of Δ*opp* to glucose or ammonium**

(A) Aggregation curve of Δ*opp* in rich medium + no glucose, 0.2% glucose, or 0.6% glucose. The top panel shows the aggregated fraction and the bottom panel shows the planktonic fraction. Data are representative of n=2 trials. (B) Aggregation curve of Δ*opp* in rich medium + no glucose with no NH_4_Cl, 25 mM NH_4_Cl, or 75 mM NH_4_Cl. The top panel shows the aggregated fraction and the bottom panel shows the planktonic fraction. Data are representative of n=2 trials.

**Figure S4 – NaCl does not affect aggregation**

Aggregation curve of WT in rich medium (no glucose) with or without 75 mM NaCl. The top panel shows the aggregated fraction and the bottom panel shows the planktonic fraction. Data are representative of n=3 trials.

**Figure S5 – Growth as planktonic cells in pyruvate defined medium with no NH_4_Cl**

Aggregation curve of WT in pyruvate defined medium (with no N_4_Cl). Data are representative of n=3 trials.

**Figure S6 – C:N regulation of aggregation/dispersal is common among NTM**

Aggregation curves of *M. abscessus* ATCC 19977, *M. fortuitum* ATCC 6841, two smooth colony *M. abscessus* subsp. *abscessus* isolates (NTM0253a and NTM0711a), and two rough colony *M. abscessus* subsp. *abscessus* isolates (NTM0253b and NTM0711b). Strains were grown in rich medium +/- 0.2% glucose (top row in each panel) and in rich medium (no glucose) +/- 75 mM NH_4_Cl (bottom row in each panel). Data are representative of at least n=2 trials.

